# Cave evolution on repeat: reuse of the same genomic regions across lineages but not across traits in *Astyanax mexicanus*

**DOI:** 10.64898/2026.01.31.703033

**Authors:** Emilie J. Richards, Rachel L. Moran, Jonathan Wiese, Morgan O’Gorman, Pierce Hutton, Aakriti Rastogi, Owen W. North, Aubrey E. Manning, Emma Y. Roback, Johanna E. Kowalko, Nicolas Rohner, Alex C. Keene, Suzanne E. McGaugh

**Affiliations:** Ecology, Evolution, and Behavior, University of Minnesota, Saint Paul, MN 55108, USA; Department of Ecology &Ecology and Evolution, University of Chicago, Chicago IL, 60637, USA; Department of Biology, Texas A&M University, College Station, TX 77843, USA; Department of Biological Sciences, Lehigh University, Bethlehem, PA 18015, USA; Institute of Integrative Cell Biology and Physiology, University of Münster, Germany

**Keywords:** Repeated evolution, Quantitative Trait Loci, Trait co-evolution, Genetic basis of sleep, cave evolution

## Abstract

Similar traits repeatedly evolve across independent populations in response to similar environmental conditions. For many repeatedly evolved traits it is unknown if populations evolve similar traits through the same or different genetic mechanisms. To address this question, we leveraged the Mexican tetra fish, *Astyanax mexicanus*, which has evolved repeatedly through altering many traits including reduced sleep duration, eye degeneration, and metabolic shifts to accommodate limited nutrient availability. We defined whether shared or independent genetic architecture govern the repeated evolution of sleep loss, increased food consumption, early onset adipose deposition, and eye loss in different evolutionary origins of the cavefish phenotype by using Quantitative Trait Locus (QTL) mapping across three cave x surface F2 mapping populations. We found that, among the traits evaluated, eye loss exhibits the most genetic repeatability, with ∼43% of QTL shared across lineages. Sleep loss and metabolic traits (i.e., feeding, adiposity) were genetically less repeatable, with only ∼25-33% of QTL shared across lineages. Next, we explored whether QTL for metabolism, eye loss, and sleep traits in cavefish co-localize in the cavefish genome and could be inherited together to facilitate potential cavefish adaptation. Although these traits have repeatedly co-evolved in cave populations, we did not find evidence for extensive genetic linkage among them. Overall, we found that genetic repeatability is a common feature in the repeated evolution of cave traits, the extent of genetic repeatability varies across cave traits, and that there is little evidence for widespread co-localization of sleep, eye loss, and metabolic traits within the genome.

**Summary:** Independent populations that evolve similar traits in response to similar environmental conditions provide us with natural replicates for studying what constrains evolutionary change. Iconic examples of repeated trait evolution often involve repeated mutations in the same genes, suggesting there are limits on the type of genes and mutations that can contribute to phenotypic evolution. Here we conduct a multi-trait, multi-population QTL analysis for similar eye loss, sleep loss and metabolic shifts across two lineages of *Astyanax mexicanus*. Genetic repeatability is present for most traits, but its extent is highly trait dependent and occurs at the level of genomic regions rather than specific genes, highlighting the mosaic of constraint and flexibility involved in genetic repeatability.

## Introduction

Repeated evolution is the shift of independent lineages to similar phenotypes in response to similar environments. Systems with repeatedly evolved phenotypes offer natural replicates to understand genetic and developmental mechanisms underlying evolutionary change (Harvey and Pagel 1991; Stern and Orgogozo 2009; Stern 2013; Storz 2016; Agrawal 2017; Langerhans 2018; Sackton and Clark 2019). These systems have been especially valuable for dissecting the genetic basis of complex traits. For example, independent freshwater populations of stickleback evolved fewer armor plates and pelvic spines through similar mutations in the same genes, *eda* and *pitx1*, respectively. Several *Heliconius* species evolved similar red patterns through independent mutations in the same enhancer elements for *optix* (Reed et al. 2011; Lewis et al. 2019; Orteu et al. 2024). Similarly, multiple *Mimulus* species have independently evolved red floral spots through mutations in the pigment forming gene *rto* (Wu et al. 2013; Ding et al. 2020). However, comparative analysis across other systems reveal that repeated trait evolution may also involve using distinct genetic paths to similar phenotypes (e.g. (Sturm and Duffy 2012; Elmer et al. 2014; Kautt et al. 2020)). This raises the fundamental question of why some case studies of repeated evolution involve reuse of the same genetic mechanisms or genomic regions while others do not.

A rich literature of theoretical work on the genetic basis of adaptation provides a foundation for exploring which how independent populations experience similar adaptive scenarios may be constrained to similar genetic pathways. For example, mutations affecting pleiotropic genes were once thought to be universally unlikely contributors to adaptation due to their potential negative fitness consequences (reviewed in (Rennison and Peichel 2022)). Yet, more recent theoretical work suggests that when adaptation occurs in the face of gene flow, pleiotropic or linked genetic architectures have a selective advantage over more disparate genetic architectures (Yeaman et al. 2016; Láruson et al. 2020; Yeaman 2022; Battlay et al. 2024). Such co-localized genetic architectures act as large-effect loci that selection can more easily drive to fixation in a segregating pool of non-adapted and adapted genotypes. Repeated use of the same loci may even occur across independent populations facing similar selection pressures to maintain local adaptation in the face of gene flow if only a few loci are capable of producing such large effect, co-localized genetic architectures (Yeaman 2022; Battlay et al. 2024). Consequently, understanding whether repeatedly evolved suites of traits share genomic regions due to pleiotropy or linkage remains central to the question of what drives repeated evolution.

The Mexican tetra fish, *Astyanax mexicanus*, provides an extraordinary opportunity to investigate these questions. Populations of *A. mexicanus* live in both surface rivers and subterranean karst landscapes in the Sierra de El Abra and Sierra de Guatemala regions of northeastern Mexico. The cave morph is found in at least 35 caves (Garduño-Sánchez et al. 2023; Miranda-Gamboa et al. 2023) and has independently evolved from at least two different surface lineages (Coghill et al. 2014; Herman et al. 2018; Moran et al. 2023). Across these independent origins, cave populations have repeatedly evolved strikingly similar phenotypes including eye and pigmentation loss, altered circadian rhythms, reduced sleep, and increased adiposity relative to surface fish (Protas et al. 2006; Duboué et al. 2011; Beale et al. 2013; Aspiras et al. 2015; Keene et al. 2015; Xiong et al. 2018; Sifuentes-Romero et al. 2020; Mack et al. 2021; Powers et al. 2023).

Additionally, many cave populations experience ongoing hybridization with surface populations (Bradic et al. 2012; Moran et al. 2022; Miranda-Gamboa et al. 2023). Similar selection pressures to maintain locally adapted suites of cave traits in the face of gene flow from surface populations could have favored the repeated retention of the same large-effect, pleiotropic, or linked loci across independently evolving populations, as theoretical predicted for (Yeaman and Otto 2011a; Yeaman and Whitlock 2011; Yeaman 2013; Yeaman et al. 2016). Past QTL and functional gene validation studies in cavefish lend empirical evidence for the existence of co-localized genetic architectures underlying multiple cave traits, from deletions in single genes like *oca2* influencing both sleep and pigmentation loss (O’Gorman et al. 2021) to the detection of several hotspots in the genome where a catalogue of diverse cave traits have been mapped to (Wiese et al. 2025). However, how repeated such genetic architectures are across independent cave populations remains unclear as genotype to phenotype studies in this system have largely been done with only a single cave population (Wiese et al. 2025). Individual cave populations do exhibit variation in these traits shifts, including differences in sleep duration, fat deposition, metabolic responses to fasting, and early eye morphology (Duboué et al. 2011; Aspiras et al. 2015; Yoshizawa et al. 2015; Jaggard et al. 2017; Xiong et al. 2018, 2022; McGaugh et al. 2020). This nuance in repeatability among traits suggests a mixture of shared and lineage-specific genetic mechanisms may be involved in the repeated evolution of cave traits.

Here, we leverage *A. mexicanus* to test the extent to which the genetic bases of sleep loss, eye loss, hyperphagia, and early-onset adiposity are repeatable across independent lineages and whether these traits co-localize in the genome. Although seemingly disparate in function, eye loss, sleep loss and metabolic shifts are likely a suite of traits that interact with each other. Sleep, metabolism and eye traits are phenotypically correlated across diverse biological systems: eye loss reduces metabolic demand, metabolic shifts influence sleep regulation, sleep and circadian rhythm genes in turn affect energy balance and eye disorders impact circadian functioning and sleep quality (Moran et al. 2014; Yasumoto et al. 2016; Lane et al. 2017; Zhou et al. 2023; Choi et al. 2025; Feeney et al. 2025). These phenotypic correlations might stem in part from shared molecular mechanisms as feeding and sleep behaviors are regulated by some of the same neuropeptides, neurocircuitry, and circadian rhythm controllers (reviewed in (Reinke and Asher 2019; Bhat et al. 2021; Shen et al. 2022; Levine et al. 2024)). These universal relationships the possibility that sleep, eye and metabolic shifts may have co-evolve through shared genetic architectures involving pleiotropy or linkage.

Using Quantitative Trait Loci (QTL) mapping in three cave x surface F2 populations we assess: 1) whether the genetic bases of a variety of traits is repeatable across independent populations, 2) whether suites of traits co-localize in the genome, 3) whether co-localization of traits in the genome is repeated across multiple populations. We find that the extent of genetic repeatability varies by trait. Additionally, we find little evidence for widespread co-localization of sleep, eye loss, and metabolic traits in the genome both within a single population or repeated across populations. Notably, however, we detect one instance shared genetic architecture among independent lineages of cavefish. Overall, this study highlights both constraint and flexibility in genomic evolution during repeated evolution to extreme environments.

## Methods

### F2 hybrid intercrosses between surface and cave populations

We leveraged the diverse phenotype combinations generated in F2 hybrid populations to assess whether sleep, metabolic, and eye phenotypes are inherited together in *A. mexicanus*. A single lab-reared surface morph female was bred with one male from three cave populations: Pachón, Tinaja, and Molino to generate three different F1 hybrid populations (Figure 1). We then crossed F1 siblings within each cross to create three F2 mapping populations. Crossing details and rearing conditions for these populations are described in the supplemental methods.

**Figure 1.**
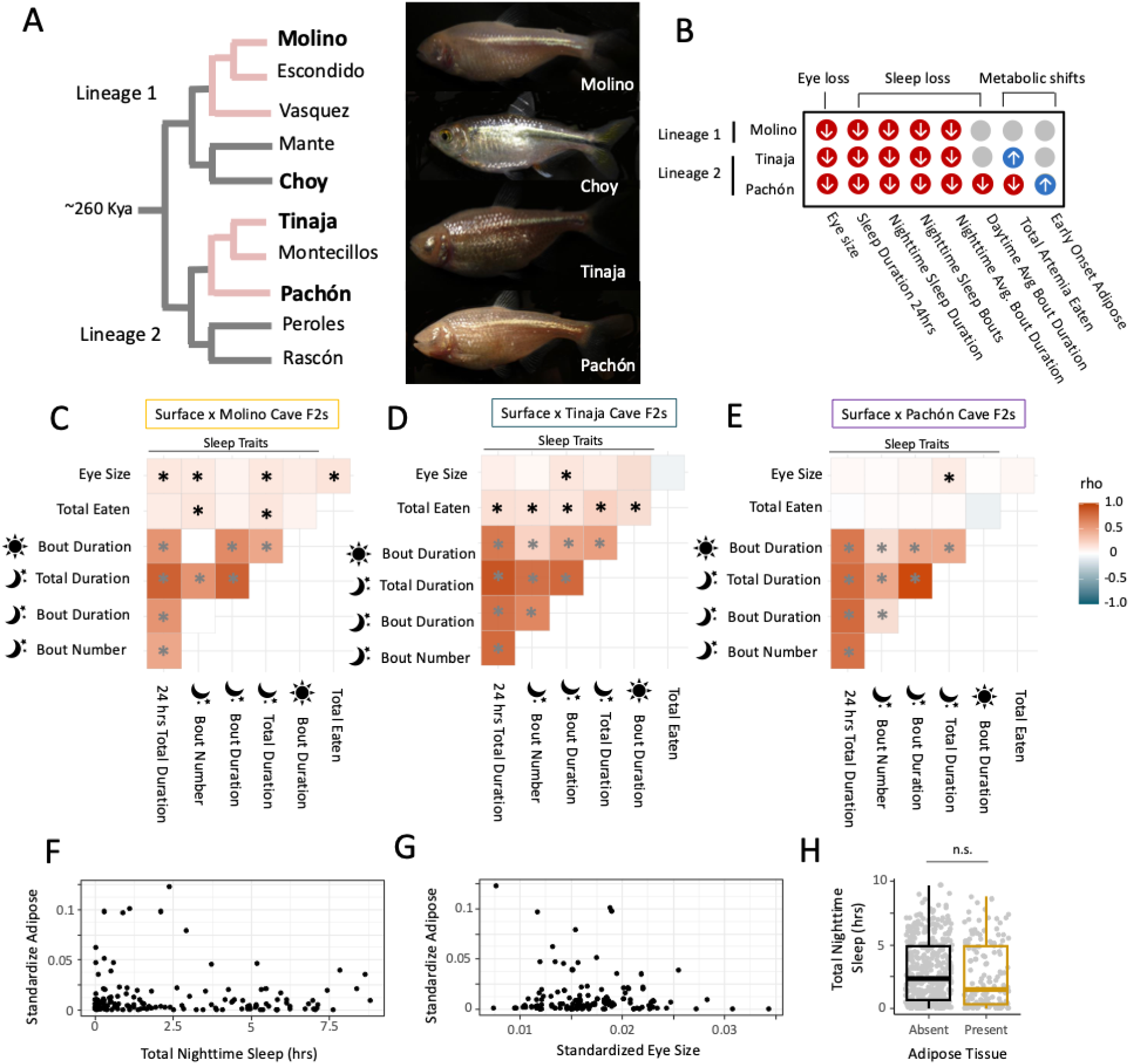
Present but weak correlations among sleep, metabolic and eye traits across independent cave lineages of *Astyanax mexicanus*. A) A cladogram with representative cave and surface populations of *A. mexicanus* highlighting the two independent evolutions of Astyanax cavefish Sierra del El Abra and Sierra de Guatemala in Northeastern Mexico. The lab populations of cave and surface fish used in this study are derived from the populations in bold. B) Focal trait shifts between surface and cavefish investigated in this study. Blue circles with an up arrow indicate that cave populations exhibit an increase in the amount of the behavior in cavefish populations compared to Río Choy surface population at 21-22 days post fertilization. Red circles with a down arrow indicate an observed decrease in the amount or size of the trait in caves compared to the Río Choy surface population at 21-22 days post fertilization. Heat map matrices of spearman’s rank correlation coefficients among cave traits in F2 hybrids between Río Choy surface and C) Molino cave, D) Tinaja cave and E) Pachón cave individuals. Lighter colors in the heat map indicates smaller Spearman’s rank correlation coefficient (rho) values and weaker relationships between the two traits states in hybrids. Asterisks indicate relationships that are significantly different from zero (*P*-value < 0.05). Black asterisks are significant relationships between different categories of cave traits (sleep, eye size, metabolism) and gray asterisks are relationships between different sleep traits. Amount of adipose tissue in surface x Pachón F2 hybrid population by 22 dpf has a complex, non-monotonic relationship with F) sleep (represented by total nighttime sleep duration) and G) eye size. Eye size and adipose tissue for an individual larval fish was standardized for size by dividing the value by the fish’s standard length. H) F2 hybrids with and without adipose tissue present slept for similar amounts of time during the night. The interquartile summary statistics of sleep for all individuals with (gold) or without (black) adipose tissue are shown in the box plot and the raw sleep values for individuals within those two groups are shown in grey circles. Only surface x Pachón F2 hybrid individuals displayed variation in adipose tissue at 22 days post fertilization (dpf).

### Parental and F2 hybrid population phenotypes

In each hybrid population and representatives of the parental populations, we measured a single eye trait, nine sleep traits, and two metabolism traits amenable to high-throughput assays in 21 days post fertilization (dpf) larval fish from the three F2 hybrid mapping populations and the four parental populations (Tables S1-S10) following the protocols described below.

#### Sleep assay

For measuring sleep in hybrid larval fish, we followed a high-throughput sleep assay previously developed for larval *A. mexicanus* (Jaggard et al. 2019). Briefly, 20 dpf larval fish were placed into individual wells of 24-well plates and filmed over a 24-hour period with a light and dark cycle. Larvae were acclimated in 24-well plates for ∼15 hours prior to recording. Following feeding with *Artemia* for 10 minutes, fish were recorded for 24 hours (14 hours of light:10 hours of dark) starting at ZT0. Sleep bouts were defined as periods of inactivity greater than 60 seconds. Inactivity was defined as periods of time where the larva’s velocity does not exceed 6 mm/s. Previous studies have shown that these periods of inactivity are strongly correlated with an increase in arousal threshold (Duboué et al. 2011; Yoshizawa et al. 2015). For each fish, we calculated the average length of each sleep bout and the total number of sleep bouts over a 24-hour period. Total sleep duration across 24 hours was the sum of the length of all sleep bouts. We additionally parsed the sleep duration between the day (the 14 hours of light) and night (the 10 hours of dark) periods to investigate shifts in circadian timing of sleep between surface and cave (Supplementary Data S1). We also measured sleep behaviors in 21 dpf representatives from the F0 populations (surface and three cave populations) following this same protocol.

#### Total eaten assay

At 21 dpf, following the 24-hour sleep recording, the feeding behavior of these fish was assayed. Fish were transferred to individual wells in a 24-well plate pre-filled with approximately 70 *Artemia* nauplii per well and recorded for two hours. Total food intake was quantified by counting any remaining *Artemia* after the feeding using the Fiji distribution of ImageJ (v1.5) and subtracting that from the total number placed in the well before the feeding trial to get the number of *Artemia* consumed in a 2-hour period (Supplementary Data S1). We also measured sleep behaviors in 21 dpf representatives from the F0 populations (surface and three cave populations) following this same protocol (Figure S1).

#### Early onset of adipose tissue assay

Following feeding, fish were fasted for 20 hours (to limit autofluorescence from any food they consumed), and at 22 dpf, fish were stained with Nile red (Sigma Aldrich 19123) to visual adipose tissue following the procedure outlined in (Xiong et al. 2018). The staining solution was prepared by dissolving Nile red in acetone at a concentration of 1.25 mg/mL and stored in the dark at −20°C. Prior to staining, the solution was diluted with fish system water to a final working concentration of 1 μg/mL. Fish were stained by placing them in a 24-well plate with 1mL of the working solution in each well. These well plates were then placed in a 28°C incubator for 30 min, as described in a previous adipose assay (O’Gorman et al. 2021). After this incubation period, fish were euthanized in 100 ug/mL MS-222 and imaged on a Nikon SMZ25 stereoscope with a GFP filter. From these images, the hybrid larval fish were scored for the presence and area of Nile red staining (Supplementary Data S1).

#### External morphology assays

Standard length and eye diameter were measured from the images described above using Fiji (v 1.5; (Schindelin et al. 2012)). Eye diameter was measured from the anterior edge of the eye to the posterior edge. Standard length was measured from the anterior tips of the snout to the base of the caudal fin. Eye diameter was corrected for standard length by dividing an individual’s eye diameter by its standard length.

#### Trait correlation analyses

As an initial assessment of which cave traits are genetically correlated, we calculated correlations between trait values among hybrid individuals. To choose an appropriate correlation statistic, we first visualized the raw relationships between different traits (e.g., eye size and total sleep duration) through scatter plots to assess whether these relationships appear linear, monotypic or non-monotypic (Figure S2-S7). Most traits had approximately linear or monotypic relationships with one another, so we calculated Spearman’s rank correlation between traits using the rcorr() function in the R package Hmisc (v5.2-3). The relationship between adipose tissue amount and sleep was non-monotypic (Figure 1), so we used a Kruskal-Wallis test to compare median sleep amount between hybrids with adipose tissue present versus absent. We used the function kruskal.test() native to base R (v4.5.0) to calculate rank sums and p-values for this assessment.

### Parents and F2 hybrid offspring genotypes

#### DNA sequencing

To determine the genotypes F2 hybrids inherited from the F0 parents, we sequenced DNA from fin tissue for each F0 parent individuals and whole-body tissue for the 22 dpf F2 individuals. For the first 384 samples, the F2 whole body tissue samples were stored in ethanol and sent to Eurofins BioDiagnostics (River Falls, WI) for DNA extractions, quantification, and normalization and then shipped to Floragenex sequence facility in Beaverton, Oregon for Restriction site Associated DNA Sequencing (RAD-Seq). RAD-Seq library creation started with a custom digest of all DNA extractions with the restriction enzyme SbfI, which we predicted would create approximately 40k cut sites in the *A. mexicanus* genome. Removal of the first 384 samples sequenced through Floragenex did not impact QTL detected downstream. Digested DNA was then prepped following the RAD-seq protocol outlined in the original single-digest RADseq protocol of (Etter et al. 2011) and sequenced using 150bp single-end Illumina HiSeq Sequencing.

The remaining 977 F2 whole body tissue samples went through DNA extractions, restriction digest, and library prep at the University of Minnesota Genomics Center using the same protocols described above. These samples were sequenced using 100bp single-end NovaSeq S1 SP sequencing at the same center.

We obtained 2.64, 1.69, and 1.73 billion total sequence reads from the 582 Surface x Pachón, 386 Surface x Molino, and 393 Surface x Tinaja F2 hybrids, respectively.

#### Raw data processing and aligning to reference genome

We first assessed read sequences for default quality metrics and removed SbfI cut site sequences from reads using the process_radtags function from Stacks (v2.54; (Catchen et al. 2011, 2013). This step checks the average quality score within windows (15% of the length of the read) and drops reads with an average raw phred scores below 10. We next used bwa-mem (v0.7.17; (Li and Durbin 2009)) to map trimmed reads to the most current version of the *A. mexicanus* surface linear reference genome (AstMex3_surface; GCA_023375975.1 (Warren et al. 2024). This reference genome has a total sequence length of 1.4 Gbp, 25 chromosomes,109 additional scaffolds and a scaffold N50 of 51.8 Mbp. We identified and removed duplicate reads using MarkDuplicates and created BAM indices using BuildBamIndex in the Picard package (http://picard.sourceforge.net<u>(v.2.0.1)</u> as required for downstream genotyping with Genome Analysis Toolkit (GATK v 3.5;(DePristo et al. 2011)).

While average read depth for an individual hybrid ranged from 1.52 to 351.6 reads per locus, the median read depth was similar across the three crosses. F2 hybrids from Surface x Molino, Surface x Pachón and Surface x Tinaja F2 cross populations were represented by a median read depth of 69.2, 73.3, and 69.9 reads per locus respectively.

#### Calling Variants

First, we called and filtered variants separately for: 1) the Surface x Pachón intercross (two F0 parents and 582 F2 hybrids), 2) the Surface x Molino (two F0 parents and 380 F2 hybrids) and 3) the Surface x Tinaja intercross (two F0 parents and 393 F2 hybrids). We used the HaplotypeCaller and GenotypeGVCFs functions from the Genome Analysis Toolkit (v 3.5;(DePristo et al. 2011)) to call and refine our single nucleotide polymorphism (SNP) variant datasets. As *A. mexicanus* lacks high-quality known allele resources, we used GATK best practice recommendations for hard filtering criteria (QD < 2.0; FS > 60; MQRankSum < -12.5; ReadPosRankSum < -8) to filter out low quality variant calls. Variants were then filtered to exclude SNPs in repetitive regions of the genome and within 3 base pairs of a GATK called indel. Repetitive regions of the genome were identified from WindowMasker files available with the reference genome on NCBI. We further filtered out genotypes with quality scores less than 20 and kept only biallelic sites using vcftools (v0.1.16; (Danecek et al. 2011)). We ended up with genotypes at 326,073 biallelic variant sites in our Surface x Pachón F2 sequence dataset,135,672 sites in our Surface x Molino F2 cross sequence dataset, and 158,326 sites in our Surface x Tinaja F2 cross sequence dataset.

We then filtered sites where many individuals were missing genotypes. We only retained sites with complete genotype information in the parents of the cross (93,933 SNPs for Surface x Pachón F2 cross; 59,075 SNPs for Surface x Molino F2 cross; 72,025 SNPs for Surface x Tinaja F2 cross). We filtered out sites that had more than 200 missing genotypes in a hybrid population, which left us with 64,311 SNP sites in the Surface x Molino F2, 76,499 in the Surface x Tinaja F2, and 102,399 in the Surface x Pachón F2 hybrid population genotype datasets at this stage

#### Selecting Variant Sites for QTL analyses

To construct linkage maps and run QTL analyses, we applied additional variant filters to our genotype datasets using functions in the R packages Rqtl (v1.46-2; (Broman et al. 2003)), and qtlTools (v1.2; https://github.com/jtlovell/qtlTools/). First, we filtered our genotype datasets down to only biallelic variant sites that were fixed for alternative alleles in the two individual F0 parents of the focal hybrid cross and then converted these vcf files into .geno files input required by Rqtl using the R package utl (v0.2.0; https://github.com/gact/utl). This reduced our Surface x Pachón F2 cross genotype dataset to 2,979 sites across 581 individuals, Surface x Molino F2 cross genotype dataset to 4,306 sites across 386 individuals, and our Surface x Tinaja F2 cross genotype dataset to 3,126 sites across 391 individuals. Next, we removed individuals with genotype information for less than 1,000 markers from downstream QTL analyses. This left us with 572 individuals from the Surface x Pachón F2 cross, 367 individuals from the Surface x Molino F2 cross, and 389 individuals from the Surface x Tinaja F2 cross to do QTL analyses with. The genotypes of hybrid individuals can be found in Supplementary Data S2.

Before constructing our genetic maps, we set aside markers that strongly deviated from predicted 1AA:2AB:1BB ratio of genotypes among F2 hybrids based on the chi-square test calculated in Rqtl function geno.table() and a Bonferroni corrected p-value < 10^-5^. This left us with 2,319, 3,664 and 2,439 variants as input into the Surface x Pachón, Surface x Molino, and Surface x Tinaja F2 cross linkage map construction. Variants were spaced apart from each other in the genome with a median distance of 0.220Mb for Surface x Pachón, 0.301Mb for Surface x Molino, and 0.398Mb for Surface x Tinaja crosses. Marker positions and the read sequences are listed in Supplementary Data S3.

### QTL Analyses

#### Marker filtration and Linkage Map Construction

We performed several quality control steps in preparation for estimating linkage maps. First, we used the rqtl function error.lod() to assign each genotype at each marker a LOD score meant to represent the likelihood that a particular genotype could be a result of genotyping error assuming an error rate across our RADseq dataset of 0.01. We removed any genotypes that had a LOD score > 4 from downstream analyses. Second, we used the rqtl function estmap() to form initial linkage groups and order markers, using the Kosambi method for calculating genetic distances and a genotype error probability of 0.05. We inspected these linkage groups for the appropriate ordering of markers based on the reference genome positions and moved any misplaced markers to their proper locations with movemarkers() function in R. Once we re-ordered markers, we re-estimated the linkage map following the same method described above. Finally, we used the dropByDropone() function in the package qtlTools (v. 1.2.0; https://github.com/jtlovell/qtlTools/) to identify potentially problematic markers that expand the linkage group by 20 cM or more when included. Small linkage groups with less than 10 markers that mapped only to unplaced scaffolds in the reference genome were dropped from the map as we did not have enough resolution to identify QTL effectively on these.

Our final genetic maps included: 1) Surface x Pachón F2 cross: 572 individuals, 2,308 markers; 25 linkage groups, and a total map length of 3550 cM; and 2) Surface x Molino F2 cross: 367 individuals, 2,656 markers, and 25 linkage groups and a total map length of 3,099 cM and 3) Surface x Tinaja F2 cross: 389 individuals, 1,884 markers, and 25 linkage groups and a total map length of 3,127 cM. Previous karyotypes of *Astyanax* species estimated 25 diploid chromosomes, matching the linkage groups in this study and chromosome-level genome assembly. The average inter-marker map distance was 1.2-1.7 cM across the three maps (Supplementary Data S3). We found 213 exact markers shared across all three linkage maps, and 399-618 exact markers shared between two of the three linkage maps (Figures S8-S12).

#### Identification of significant QTL peaks

We mapped QTL for eight traits: total sleep duration across 24 hours, total sleep duration during night, total number of sleep bouts during night, average length of sleep bouts during night and average length of sleep bouts during day, total *Artemia* eaten, eye size and early onset of adiposity. We selected these eight traits because they were significantly different in size or amount between at least one surface and one cave parent populations at 21 dpf from a mixed model we ran using lmer function in the R package lme4 (v1.1-32;(Bates et al. 2015)). The mixed models included population and day period (e.g., day or night) if applicable as fixed effects and individual ID and well position in the assay as random effects (Tables S1-10).

Since all three crosses shared a F0 surface female, we used a joint mapping approach that combined all hybrid individuals to increase power to detect any small effect QTL that were common across mapping populations. Joint mapping and QTL scans were done using statgenMPP (v1.0.2; (Li et al. 2022)). This approach integrates identity-by-descent calculations for offspring inheriting a particular parent’s genetic variation with linear mixed models to estimate random QTL effects relative to genetic ancestry while considering that polygenic and family background genetic variation is present. This method requires a single linkage map for the combined hybrid populations, so we created a consensus map of the three individual hybrid population linkage maps using R package LPmerge (v1.7;(Endelman and Plomion 2014). Comparisons of marker positions and density across the consensus linkage map and independently assembled linkage maps for each population are provided in the supplementary materials (Table S12; Figure S1-S5). We then ran a joint mapping QTL scan for the seven traits that were variable in all three hybrid populations. Early on-set of adipose tissue by 22 dpf was only variable among Surface x Pachón F2 hybrids and was excluded from the joint mapping analysis.

For the joint mapping analysis, we scanned the combined linkage map and genetic markers for the three hybrid populations for associations between hybrid genotypes and phenotypes by calculating LOD scores in 10 cM windows across the genome. A significant association between a 10cM region and a phenotype was set at a threshold LOD score with a negative log_10_ transformed P-value higher than 2. This threshold was determined from a Bonferroni correction with an alpha threshold of 0.10 adjusted for the number of markers in the map. For adipose tissue, we conducted a single population, multiQTL scan with only the genotype, phenotype, and linkage map data of the Surface x Pachón F2 hybrid population. We ran this QTL analysis with the logistic regression model option ‘binary’ in the scan1() function of R/qtl2 package. To find QTL peaks, we used a permutation-based 10% significance threshold for LOD scores calculated with the scan1perm() with 1000 permutations and the model argument set to ‘binary’.

### Assessment of genetic repeatability in the repeated evolution of cave traits

#### Colocalization of QTL for sleep, metabolic and eye traits

To determine if the phenotypic correlations observed in the F2 hybrids are the result of shared genetic architecture among traits, we assessed whether any QTL from across two or more of the four trait categories (sleep, eye size, total eaten, and adipose tissue) overlapped. Overlapping QTL regions were defined as 1) having the same estimated peak markers and boundary markers or 2) the estimated boundaries of a detected QTL for one trait was nested within the boundaries of another. Varying the allowed distance at which multiple independent QTL could be detected between (e.g. two QTLs for a trait could not be called within 10 cM or 20 cM of each other) did not change the assessment of colocalization across independent lineages except in the case of two consecutive QTL on chromosome 5 associated with sleep loss (this exception is discussed in the supplementary materials; Table S13).

#### Repeated use of the same genomic regions across independent caves

We also assessed whether genetic inheritance from all three cave parents contributed to genotype and phenotype associations in QTL detected from the combined dataset. We did this using a ‘drop one’ strategy where we removed one population at a time from the combined dataset and reran the statgenMPP scan. QTLs that were not detected when the genotypes and phenotypes of a particular cross was left out of the QTL scan indicated that the cave parents’ ancestry contribute significantly to the phenotype-genotype association in that region. Next, we qualitatively compared the effect size estimates that statgenMPP calculates for the genotypes of the individual parents on QTL (Tables S14-S16). statgenMPP calculates effect estimates of parental genotypes on hybrid phenotypes relative to the average phenotype of all hybrids. Parental genotypes in QTL regions with estimated effect values near zero indicate that the ancestry from this parent does not contribute to the phenotype-genotype association. Lastly, we also explored several alternative QTL analyses including independent mapping across each hybrid population under both single and multi-QTL models and present those results and comparisons across all QTL approaches in the supplementary materials (Supplementary Methods, Tables S17-S19). The improved power of the joint estimate approach is notable, as the single population approach detected fewer shared QTL peaks across populations.

#### Repeated selection on the same genes in shared QTL regions

Next, we assess whether shared QTL signatures were driven by repeated selection on the same set of genes. To do this, we leveraged a recently published selective sweep dataset estimated from the genomes of wild-caught cave and surface populations (Roback et al. 2025). The individuals used in this dataset derive from the same parent populations that the QTL cross parental generations do (Molino cave n=14, Pachón cave n=19, and Tinaja cave n=21, Río Choy surface n=9); as well as the wild Rascón surface population (n=13) to represent Lineage 2 surface fish. These genomes were mapped to the same reference genome that we mapped the F2 hybrid individuals to for the QTL analysis (GCA_023375975.1). Selective sweeps were identified using diploS/HIC (v1.0.6;(Kern and Schrider 2018)), a neural network approach that uses simulated genetic variation under different selective and neutral scenarios to train a model to identify selective sweep patterns in real populations. Training sets of genetic variation for cave and surface populations included demographic parameters to account for historical changes in population sizes and reduce false positives of sweeps in regions of the genome that experience drastic but random changes in genetic variation. These demographic parameters are the same used in Moran et al. (2023). With this model, 5-kb windows across the genome were classified as either evolved neutrally, experienced a hard sweep or soft sweep.

Next, we sorted 5-kb windows in this dataset into two groups: windows with evidence of selection in the form of a selective sweep and windows that are neutrally evolving. We did not distinguish between hard and soft selective sweeps in this study, as validating the distinction between the two types of sweeps is challenging (Kern and Schrider 2018). The colonization of cave environments likely involved strong bottlenecks that could have swept variants randomly to high frequency in resulting cave populations, which are difficult to distinguish from variants swept to high frequency by selective forces. We further filtered selective sweeps to those that were robust to possible model misspecification in parameters like bottleneck severity by only retaining calls that were consistent between runs where we applied either the cave demographic model or the surface demographic model to a cave population. Although this filter is conservative and likely discards regions with complex signatures of selection from downstream analyses, this allowed us to focus on evidence of genetic repeatability driven by similar selective sweeps across cave populations with variable but unknown demographic histories that could also influence fixation. We applied a similar model misspecification robustness filter to the surface populations. Lastly, we looked for selective sweep signatures likely to be driven by independent, repeated mutations among cave populations to assess whether genetic repeatability detected from the QTL analysis is driven by independent but repeated selective sweep events. We parsed out genes that experienced selective sweeps in cave populations from both lineages (Molino and Pachón or Tinaja) but not in surface populations (Río Choy and Rascón). A synthesized list of selection signatures that overlapped with coding regions in all genes within QTL regions along with gene annotations from model organisms and a summarized output of the diploS/HIC results across all genes in the genome are provided in Supplementary Data S4 and S5.

#### Validation of new candidate gene for sleep regulation in Astyanax mexicanus

To illustrate the utility of the QTL experiments in *A. mexicanus* for understanding the genetic basis of complex traits such as sleep, we selected a gene candidate residing in a sleep QTL region to functionally validate. We created crispant surface fish harboring mutations in *tgfa*: a gene that fell within the sleep and eye size QTL region on chromosome 5 and has recently been shown in zebrafish to reduce sleep when functionally disrupted (Lee et al. 2019). Briefly, we assayed 22 dpf surface fish injected with Cas9 protein and a gRNA targeting *tgfa*, along with their wild-type, non-injected siblings, for sleep behavior over a 24-hour period. Details on how crispants were generated and their sleep patterns were assayed can be found in the supplementary methods S6. We used a Kruskal-Wallis test to assess significant differences in the median amount of sleep bout duration, number of sleep bouts, and total sleep duration across the 24-hour trial using the kruskal.test() function in base R (v4.5.0).

## Results

### Repeated yet weak correlations among phenotypes within populations

We compared eye, sleep, and metabolic phenotypes among 21 dpf larval fish from the three cave and one surface parent populations to determine what cave phenotypes are repeated among the independent lineages by early developmental stages. Eight of the traits we measured were significantly different between one or more cave and surface populations at 21 dpf (Figure 1B & S6; Tables S3-S10): eye size, total sleep duration across 24 hours, average sleep bout duration across 24 hours, average sleep bout duration during the night, average sleep bout duration during the day, total number of nighttime sleep bouts, nighttime total sleep duration, and total *Artemia* nauplii eaten.

To gauge whether cave sleep, metabolic, and eye phenotypes are genetically correlated in *A. mexicanus* populations, we looked for correlations between phenotypes in the F2 hybrid individuals. The sleep metrics (e.g. total sleep duration and average sleep bout duration) were the most strongly correlated with each other of all the traits based on Spearman’s rank tests (Figures S7-S12). F2 hybrid populations varied widely in whether eyes, metabolic and sleep traits were correlated with each other though. Total sleep duration had a weakly positive but significant correlation with 1) total *Artemia* nauplii eaten in hybrid populations (Surface x Molino and Surface x Tinaja F2s; Figure 1C-E; Table S20) and 2) eye size (Surface x Molino and Surface x Pachón F2s; Figure 1C-E; Table S20), suggesting potential genetic correlations. Eye size and total amount of *Artemia* nauplii eaten had a weakly positive but significant relationship with each other, but only in Surface x Molino F2 hybrids (Figure 1C; Table S20). Surface x Pachón F2 hybrids with and without adipose present at this developmental stage didn’t sleep significantly different amounts from each other (Figure 1H). While population-dependent, the weak positive correlations among eye size, sleep, and total eaten traits in hybrids suggests that they are partially correlated at the genetic level.

### Sleep, feeding, and eye development have a highly polygenic architectures in cavefish

Next, we leveraged the shared female surface grandparent among the three hybrid crosses to combine the genotype and phenotype data of all hybrids into one QTL scan to find the genetic bases of the repeated traits among cave populations. The joint mapping approach detected 48 QTL for the focal traits that were phenotypically variable among all three hybrid populations (Figure 1B;Tables 1-3). For the early onset of adipose tissue, the single population multiQTL scan detected only one significant QTL (Figure 2; Tables 1-3). These QTL are found in 31 distinct QTL regions across 19 of the 25 chromosomes in the genome based on overlapping confidence intervals between the individual QTLs (Tables 1-3 & S14-16). There were 15 distinct QTL regions associated with eye size in this system, 10 associated only with sleep loss, three associated only with total *Artemia* eaten, and one associated only with early onset of adiposity (Figure 2, Tables 1-3).

**Figure 2.**
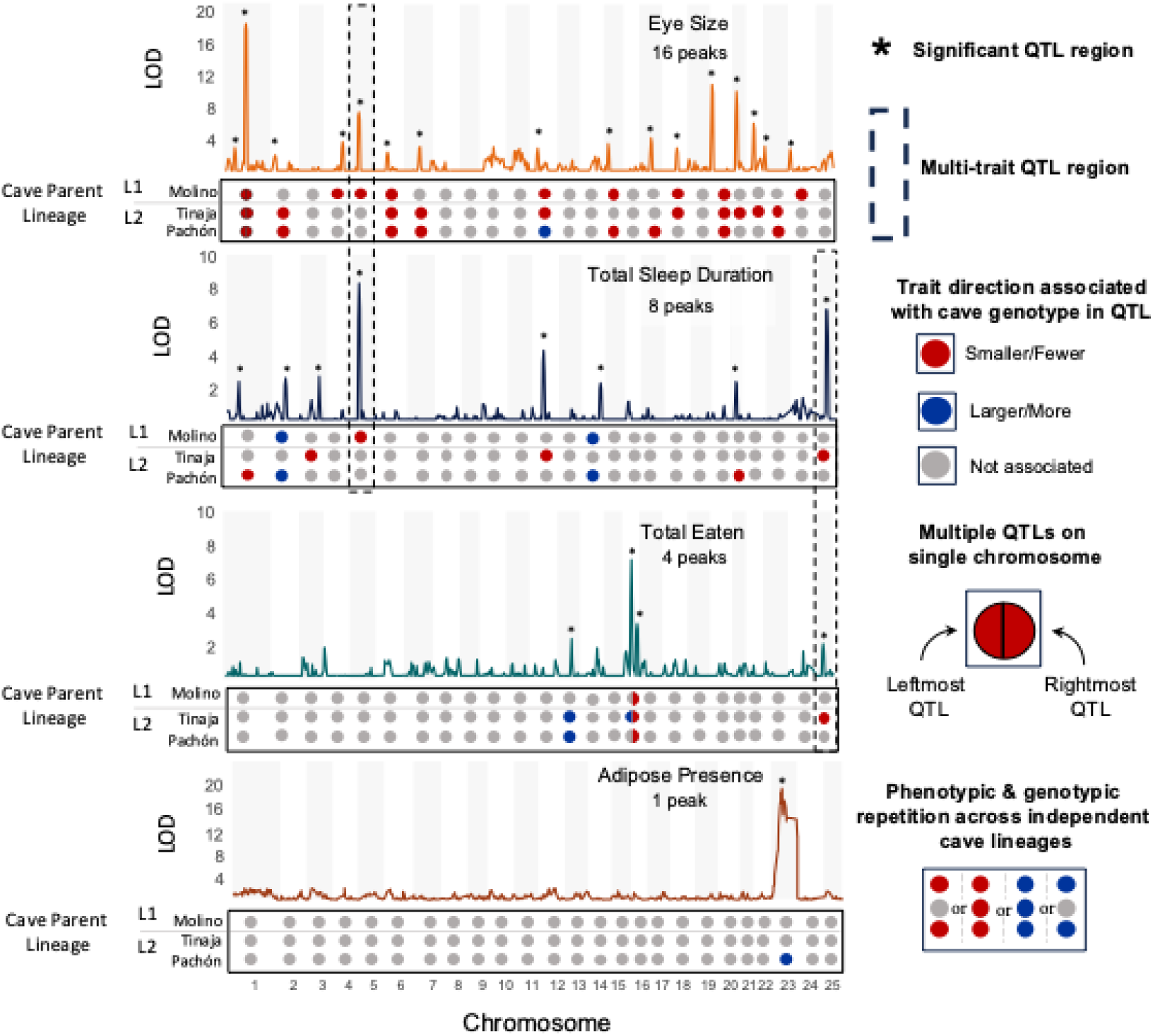
QTL scans associating genome regions with cave traits in larval fish from three surface x cave F2 hybrid populations. In this overview figure, total sleep duration is used as the representative trait for sleep loss. QTL for the other sleep metrics are featured in Figure S14. LOD scores represent the strength of association between genotypes and the four broad phenotypes of interest calculated from joint association mapping across all three surface x cave F2 hybrid crosses that share a common surface parent. The trait each panel represents is indicated in the upper right corner alongside the number of regions that were significantly associated with the traits of interest (peaks represented by asterisks). Dash-line boxes highlight genome regions in which multiple traits (e.g. eye size and sleep duration) map to the same place. Upset plots below each panel indicate which cave parent ancestry underlies the association between genotype and phenotype detected and the color of the circle represents the directionality of the association. QTL peaks in which the cave allele at the peak marker is associated with larger trait value than surface ancestry is indicated by a blue circle, and QTL peaks in which the cave ancestry is associated with a smaller trait value than surface ancestry is indicated by a red circle. Each chromosome is represented by a single circle and half colored circles are used to show phenotypic direction associated with cave ancestry when two QTL peaks are present on a single chromosome.

**Table 1.** Genomic regions detected as significantly associated with traits of interest from a genome-wide multiple QTL joint interval mapping scan. The estimated interval (chromosome:start-end) and LOD score indicating the strength of association for each QTL. The QTL region that one or more overlapping QTL were detected in are labelled ES1-ES16 and shaded in alternate colors of grey and white. Any overlapping QTL within the study where a different trait also mapped to the same genomic region are represented with acronyms (e.g. S4 from Table 2 overlaps with region ES5). Many of the QTL detected in this study were also independently detected in other QTL studies on related traits that mostly used Pachón cavefish. The QTL studies where similar traits map to an overlapping region with our QTL are referenced with superscripts. Traits from previous QTL studies that mapped to the same region but were not related to the eye size are indicated as ‘Other cave traits.’ The specific trait QTL studies our eye size QTL overlap with are listed in detail in Table S30. Phenotypic direction that cave parent ancestry was associated with are represented by arrows next to the cave parent names with downward arrow indicating a smaller average trait value than individuals with surface parent ancestry at the QTL locus and upward arrow indicating a larger average trait value. Bold parallel QTL are those in which the cave parent alleles had effects in the same phenotypic direction across multiple caves (e.g., smaller eyes). The effect size estimates for each QTL can be found in Table S13.

| QTL ID | Region (Mbp) | LOD | Overlap within study | Overlap with other QTL studies | Parallel across lineages QTL | Lineage Specific QTL |
| --- | --- | --- | --- | --- | --- | --- |
| ES1 | chr1:12.9-20.8 | 3.15 | - | Eye Size<sup>1,2</sup>,<br>Other cave traits | **Tinaja<sup>↓</sup>, Pachón<sup>↓</sup>, Molino<sup>↓</sup>** | - |
| ES2 | chr1:50.4-58.2 | 18.54 | - | Eye Size<sup>1,2</sup>,<br>Other cave traits | **Tinaja<sup>↓</sup>, Pachón<sup>↓</sup>, Molino<sup>↓</sup>** | - |
| ES3 | chr2:2.57-3.99 | 2.19 | - | Other cave traits | - | **Tinaja<sup>↓</sup>, Pachón<sup>↓</sup>** |
| ES4 | chr4:48.2-51.9 | 3.86 | - | Other cave traits | - | Molino<sup>↓</sup> |
| ES5 | chr5:12.8-19.3 | 7.54 | S4 | Eye Size<sup>1,3</sup>,<br>Other cave traits | - | Molino<sup>↓</sup> |
| ES6 | chr6:8.86-14.8 | 2.59 | - | Other cave traits | **Tinaja<sup>↓</sup>, Pachón<sup>↓</sup>, Molino<sup>↓</sup>** | - |
| ES7 | chr7:27.7-30.9 | 3.24 | - | Eye Size<sup>4</sup>,<br>Other cave traits | - | **Tinaja<sup>↓</sup>, Pachón<sup>↓</sup>** |
| ES8 | chr12:3.62-5.41 | 3.06 | - |  | **Tinaja<sup>↓</sup>, Pachón<sup>↑</sup>, Molino<sup>↓</sup>** | - |
| ES9 | chr15:10.8-14.9 | 3.61 | - | Other cave traits | **Pachón<sup>↓</sup>, Molino<sup>↓</sup>** | - |
| ES10 | chr17:12.3-18.9 | 4.36 | - | Eye Size<sup>1,3</sup>,<br>Other cave traits | - | Pachón<sup>↓</sup> |
| ES11 | chr18:23.9-29.2 | 3.09 | - | Other cave traits | **Tinaja<sup>↓</sup>, Molino<sup>↓</sup>** | - |
| ES12 | chr20:0.86-1.5 | 10.98 | - | Eye Size<sup>1,3,5</sup> | **Tinaja<sup>↓</sup>, Pachón<sup>↓</sup>, Molino<sup>↓</sup>** | - |
| ES13 | chr21:18.1-21.3 | 10.13 | - | Eye Size<sup>1,2</sup> | - | Tinaja<sup>↓</sup> |
| ES14 | chr22:18.9-26.4 | 6.16 | - | Eye Size<sup>6</sup> | - | Tinaja<sup>↓</sup> |
| ES15 | chr23:0-2.08 | 3.30 | - | - |  | **Tinaja<sup>↓</sup>, Pachón<sup>↓</sup>** |
| ES16 | chr24:0.37-1.85 | 2.95 | - | - | - | Molino<sup>↓</sup> |

<sup>1</sup>(Protas et al. 2007) <sup>2</sup>(Protas et al. 2008) <sup>3</sup>(O'Quin and McGaugh 2015) <sup>4</sup>(Yoshizawa et al. 2012) <sup>5</sup>(Powers et al. 2020) <sup>6</sup>(Kowalko et al. 2013)

### Genetic architectures repeated across independent cave populations

Cave trait evolution in both cave lineages, which represent different colonization events of cave environments by at least independent surface lineages, involves repeated divergence in some of the same genomic regions. The extent of this genetic repeatability was variable but present among all the repeated traits we examined (excluding the non-repeated early on-set of adipose tissue).

Genetic repeatability was highest for repeated eye loss among independent cave populations. Nine of the 16 (43%) eye size QTLs regions were associated with cave genotypes from both Lineage 1 (Molino) and Lineage 2 (Tinaja and/or Pachón). The remaining eye size QTL were unique to specific populations or shared only between Lineage 2 populations (Pachón and Tinaja). Notably, some Molino-specific eye size QTL (ES4, ES5) overlap in confidence intervals with eye size QTLs from previous Pachón x Surface F2 mapping studies (Table 1). Thus, up to 56% of the eye size QTL are regions of the genome that are likely repeatedly involved in the independent loss of eyes across cavefish populations.

Sleep and metabolic traits also contain repeated aspects of their genetic architectures across the two independent lineages. We found 33% of the 12 sleep QTL regions (Table 2 and S14) were shared across the independent lineages, 25% of QTL regions were shared only among Lineage 2 populations (Pachón and Tinaja), and 42% of QTL regions were unique to individual populations. For total *Artemia* eaten, only one QTL was shared across the independent lineages (Table 3; Figure 2) and one was shared within Lineage 2 (Pachón and Tinaja). The remaining two QTLs for total *Artemia* eaten (TE) were unique to Tinaja, the only population that appears to be significantly more hyperphagic than the surface population at this development stage.

**Table 2.** Genomic regions detected as significantly associated with traits of interest from a genome-wide multiple QTL joint interval mapping scan. The estimated interval (chromosome:start-end) and LOD score indicating the strength of association for each QTL. All QTL detected for sleep traits are represented by the following acronyms: TSD: total sleep duration across 24 hours; TSDN: total sleep duration at night; ASD: average durations of sleep bouts during day; ASN: average duration of sleep bouts during night; and BNN: total number of sleep bouts during night). Independent QTL regions where one or more QTL across the multiple sleep traits were detected were give the IDs S1-S12 and are shaded in alternate colors of grey and white. Multiple sleep QTL that mapped to same region of the genome are grouped together under the same QTL Region ID. Many of the QTL detected in this study were also independently detected in other QTL studies in *Astyanax mexicanus* for related traits. Phenotypic direction that cave parent alleles were associated with are represented by arrows next to the cave parent names with downward arrow indicating a smaller average trait value than individuals with surface parent ancestry at the QTL locus and upward arrow indicating a larger average trait value. Bold QTL are those in which the cave parent alleles had effects in the same phenotypic direction across multiple caves (e.g., sleeps less). All QTL peaks detected amongst the individual sleep trait scans are listed here, although several peaks map to overlapping regions on the same chromosome and are non-independent of each other given their nested nature (e.g., sleep bout number contributes to total sleep duration). Non-independent QTL peaks are collapsed and counted as one independent QTL region throughout the text. The effect size estimates for each QTL can be found in Table S13. Regions with significant QTL peaks from previous QTL studies are indicated with ‘other cave traits’. The specific trait QTL studies our sleep QTL overlap with are listed in detail in Table S31. Genes names are included where we have functional evidence from *A. mexicanus* null mutants that a gene residing in a QTL impacts sleep patterns.

| QTL Region ID | Trait | QTL ID | Region (Mbp) | LOD | Overlap within study | Overlap with other QTL studies | Shared across Lineages QTL | Lineage Specific QTL |
| --- | --- | --- | --- | --- | --- | --- | --- | --- |
| S1 | Total Sleep Duration | TS1 | chr1:24.1-29.8 | 2.55 | - | Other cave traits | - | Pachón<sup>↓</sup> |
|  | Avg. Sleep Bout Duration at Night | ASN1 | chr1:24.1-29.8 | 2.98 | - | Other cave traits | - | Pachón<sup>↓</sup> |
|  | Avg. Sleep Bout Duration Day | ASD1 | chr1:24.1-29.8 | 2.04 | - | Other cave traits | - | Pachón<sup>↓</sup> |
| S2 | Total Sleep Duration | TS2 | chr2:18.5-23.8 | 2.75 | - | Other cave traits | **Pachón<sup>↑</sup>, Molino<sup>↑</sup>** | - |
|  | Total Sleep Duration at Night | TSN1 | chr2:18.5-23.8 | 2.92 | - | Other cave traits | **Pachón<sup>↑</sup>, Molino<sup>↑</sup>** | - |
|  | Bout Number at Night | BNN1 | chr2:18.5-23.8 | 2.72 | - | Other cave traits | **Pachón<sup>↑</sup>, Molino<sup>↑</sup>** | - |
| S3 | Total Sleep Duration | TS3 | chr3:45.6-55.4 | 2.84 | - | Other cave traits | - | Tinaja<sup>↓</sup> |
|  | Bout Number at Night | BNN2 | chr3:45.7-55.4 | 2.15 | - | Other cave traits | - | Tinaja<sup>↓</sup> |
|  | Avg. Sleep Bout Duration Day | ASD2 | chr3:45.6-55.4 | 2.09 | - | Other cave traits | - | Tinaja<sup>↓</sup> |
| S4 | Total Sleep | TS4 | chr5:7.97-12.9 | 8.39 | - | Other cave traits | - | Molino<sup>↓</sup> |
|  | Avg. Sleep Bout Duration Day | ASD3 | chr5: 7.97-12.9 | 3.58 | - | Other cave traits | - | Molino<sup>↓</sup> |
|  | Total Sleep Night | TSN2 | chr5:10.4-15.8 | 3.18 | ES5 | Other cave traits | - | Molino<sup>↓</sup> |
|  | Avg. Sleep Bout Duration Night | ASN2 | chr5:10.4-15.8 | 4.87 | ES5 | Other cave traits | - | Molino<sup>↓</sup> |
|  | Total Sleep Night | TSN3 | chr5:15.8-21.2 | 2.02 | ES5 | Other cave traits<br>*oca2*<sup>1</sup> | - | **Tinaja<sup>↓</sup>, Pachón<sup>↓</sup>** |
| S5 | Total Sleep | TS5 | chr12:7.97-11.7 | 4.40 | - | Other cave traits | - | Tinaja<sup>↓</sup> |
|  | Total Sleep Night | TSN4 | chr12:7.97-11.7 | 2.93 | - | Other cave traits | - | Tinaja<sup>↓</sup> |
|  | Avg. Sleep Bout Duration Night | ASN3 | chr12:6.5-8.8 | 2.01 | - | Other cave traits,<br>*rorca*<sup>2</sup> | - | **Tinaja<sup>↓</sup>, Pachón<sup>↓</sup>** |
| S6 | Total Sleep | TS6 | chr14:40.8-43.8 | 2.46 | - | Other cave traits | - | **Tinaja<sup>↑</sup>, Pachón<sup>↑</sup>** |
| S7 | Avg. Sleep Bout Duration Day | ASD4 | chr16:28.4-35.9 | 2.27 | - | Other cave traits | **Pachón<sup>↓</sup>, Molino<sup>↓</sup>** | - |
| S8 | Avg. Sleep Bout Duration Day | ASD5 | chr17:4.8-7.8 | 2.12 | - | Other cave traits | - | Pachón<sup>↑</sup> |
| S9 | Total Sleep Night | TSN5 | chr20:18.4-24.4 | 2.12 | - | Other cave traits | - | **Tinaja<sup>↑</sup>, Pachón<sup>↑</sup>** |
| S10 | Total Sleep | TS7 | chr21:0-7.3 | 2.51 | - | - | - | Pachón<sup>↓</sup> |
|  | Total Sleep Night | TSN6 | chr21:0-7.3 | 2.68 | - | - | - | **Tinaja<sup>↓</sup>, Pachón<sup>↓</sup>** |
|  | Bout Number Night | BNN3 | chr21:0-7.3 | 2.61 | - | - | **Tinaja<sup>↓</sup>, Molino<sup>↓</sup>** | - |
| S11 | Total Sleep | TS8 | chr25:6.07-7.31 | 6.82 | TE4 | Other cave traits | - | Tinaja<sup>↓</sup> |
|  | Total Sleep Night | TSN7 | chr25:6.07-7.31 | 3.61 | TE4 | Other cave traits | - | Tinaja<sup>↓</sup> |
|  | Bout Number Night | BNN4 | chr25:4.2-6.3 | 4.09 | TE4 | Other cave traits | - | Tinaja<sup>↓</sup> |
| S12 | Avg. Sleep Duration Day | ASD6 | chr25:14-17.3 | 6.15 | - | - | - | Tinaja<sup>↓</sup> |

<sup>1</sup>(O’Gorman et al. 2021) <sup>2</sup>(Mack et al 2021)

**Table 3.** Genomic regions detected as significantly associated with traits of interest from a genome-wide multiple QTL joint interval mapping scan. The estimated interval (chromosome:start-end) and LOD score indicating the strength of association for each QTL. Any overlapping QTL where multiple traits map are represented with acronyms (TE: Total *Artemia* eaten in 2-hour period). Phenotypic direction that cave parent alleles were associated with are represented by arrows next to the cave parent names with downward arrow indicating a smaller average trait value than individuals with surface parent ancestry at the QTL locus and upward arrow indicating a larger average trait value. Bold parallel QTL are those in which the cave parent alleles had effects in the same phenotypic direction across multiple caves (e.g., eats less). Non-independent QTL peaks are collapsed and counted as one independent QTL region throughout the text. The effect size estimates for each QTL can be found in Table S13. Regions where significant QTL peaks in previous QTL studies also mapped to are indicated with ‘other cave traits’. The specific trait QTL studies our metabolism-related QTL overlap with are listed in detail in Table S32.

| Trait Category | Trait | QTL ID | Region (Mbp) | LOD | Overlap within study | Overlap with other QTL studies | Parallel across Lineages QTL | Lineage specific QTL |
| --- | --- | --- | --- | --- | --- | --- | --- | --- |
| Eating Behavior | Total *Artemia* Eaten | TE1 | chr13:34.4-40.2 | 2.48 | - | Other cave traits | - | **Tinaja<sup>↑</sup>, Pachón<sup>↑</sup>** |
|  |  | TE2 | chr16:15.1-25.1 | 7.11 | - | Other cave traits | - | Tinaja<sup>↑</sup> |
|  |  | TE3 | chr16:37.7-39.7 | 3.36 | - | Other cave traits | **Tinaja<sup>↓</sup>, Pachón<sup>↓</sup>, Molino<sup>↓</sup>** | - |
|  |  | TE4 | chr25:4.19-6.26 | 2.17 | S11 | Other cave traits | - | Tinaja<sup>↓</sup> |
| Adiposity | Presence of Adipose Tissue | EOA1 | chr23:9.61-13.8 | 20.11 | - | Other cave traits | - | Pachón<sup>↑</sup> |

### Few cave-unique and repeated signatures of selective sweeps on genes within shared QTL regions

To investigate whether the shared QTL we detected come from repeated, strong selection across the independent cave lineages on the same set of genes, we ran selective sweep scans across the genomes of wild-caught cave and surface individuals of the parental populations. Out of all 2,433 genes inside QTL regions, only two genes had signatures of selection in all three wild cave parent populations, but not in surface populations (Supplementary Data S4). Both genes were found in single trait QTL regions unique to one cave lineage (*klhl43* in Sleep QTL S3 and *zbtb17* in Total Eaten QTL TE1; Tables S21 & S22). When we widen the search to genes that were uniquely under selection in at least one cave population from both lineages (e.g. Molino and either Pachón or Tinaja), the tally increased to 30 genes (Supplementary Data S4). Eleven of the 30 genes are found within three eye size QTL and two sleep QTL shared across lineages, while the rest are found within single lineage QTL regions. This pattern does not appear to be driven by a lack of cave-unique signatures of selection in genes inside these QTL regions. Only two of the 31 QTL regions did not contain any genes with cave-unique selective sweeps (Table S23). In the remaining 29 QTL regions, 7% to 47% of the protein-coding genes were under selection in a cave population but not in either of the surface populations (Table S23). The lack of repeated, cave-specific selection signatures inside shared eye, sleep and total eaten QTL regions suggests that genetic repeatability for these traits is likely not constrained by the need for rare, singularly advantageous mutations arising in select large-effect loci.

### Limited shared genetic architecture between repeatedly coevolving cave traits

We detected only two QTL regions where multiple traits exhibited overlapping QTL intervals: one region of chromosome 25 associated with both sleep loss and total eaten and one region of chromosome 5 associated with both eye and sleep loss (Figure 2, Tables 1-3). The chromosome 25 QTL is associated with shorter total sleep duration and fewer sleep bouts during the night and less *Artemia* eaten (Figure 2, Table 2-3). Only Tinaja cave parent ancestry in the chromosome 25 QTL region is strongly associated with these phenotypes (Figure 2, Table 2-3). Residing in this region are two interesting candidate genes for sleep and metabolism regulations: *acsl1a* and *mtnr1aa* (Supplementary Data S4). Only Río Choy and Rascón surface populations have experienced a selective sweep on variation inside *acsl1a*, while Molino, Tinaja, Río Choy and Rascón populations have experienced a selective sweep on variation inside *mtnr1aa*.

For chromosome 5, sleep duration was associated with genotypes from all three cave populations but only Lineage 1 Molino cave ancestry was also associated with eye size. Previous work also mapped an eye size QTL for surface x Pachón F2 hybrids to this region, suggesting this region is associated with eye size in Lineage 2 cavefish populations as well (reviewed in Wiese et al 2025). Several interesting candidate genes with the potential to regulate or interact with sleep occur in this region, including *oca2, tgfa, ube3a*, and *ghrl* (Supplementary Data S4). *oca2* is functionally validated for impacts on sleep, pigmentation, vision-dependent prey capture behavior and stress behavior in *A. mexicanus* (Protas et al. 2006; Klaassen et al. 2018; O’Gorman et al. 2021; Choy et al. 2025b,a). In this study, we validate that genetic variation in *tgfa* can also modify sleep patterns in *A. mexicanus*. The *tgfa* crispant surface fish had a median 36% reduction in sleep compared to their wildtype siblings in a 24-hour sleep trial (Fig S14; Supplementary Data S6). The median duration of sleep bouts in *tgfa* crispants was significantly shorter than wildtype counterparts, while the total number of bouts were similar between the two groups (Fig S14; Table S29; Supplementary Data S4).

The QTL for the early onset of adipose tissue in Pachón F2 hybrids did not overlap any of the three other types of traits examined (Figure 2; Table 3), though, an eye size QTL is found ∼ 6 MB upstream of the adipose QTL (Figure 2, Table 1). Two interesting candidate genes for regulating adipose tissue that reside in this QTL are *cepba* and *arhgap1* (Supplementary Data S4).

In sum, we find that eye size, sleep amount, total amount eaten and adipose tissue map to regions of the genome that house many promising candidate genes with relevant annotations in model organisms for cave phenotypes. However, we find limited evidence that the traits of eye development, sleep, feeding, or adipose deposition are extensively genetically co-localized.

## Discussion

Dissecting the phenotypic and genotypic variation underlying repeated evolution of complex traits continues to reveal striking levels of genetic reuse in nature(Wortel et al. 2023). Our study provides an unparalleled resolution of genomic signatures associated with cave evolution, identifying both repeated and lineage-specific genetic architectures that contribute to shifts in eye size, sleep, feeding, and metabolic function. Collectively, our findings show that: (1) the degree of genetic repeatability across different cave populations varies across traits, with eye size showing the most consistent genetic basis across lineages; (2) sleep and metabolic traits exhibit more population-specific architectures; and (3) co-localization of QTL for multiple traits within the same genomic regions is rare in our data, suggesting that pleiotropy or linkage are generally not impacting the cave-derived traits examined here.

### Extensive genetic repeatability despite variable phenotypic repeatability

Cave populations of *A. mexicanus* display substantial phenotypic divergence from surface forms, but the magnitude and direction of these changes vary across caves. Population-specific differences are evident in early eye development (Sifuentes-Romero et al. 2020), pigmentation loss (Protas et al. 2006; Gross et al. 2009), sleep loss (Duboué et al. 2011), increased adiposity, and hyperphagy (Aspiras et al. 2015). For example, there is abundant intercave variation in the extent to which sleep phenotypes differ from surface populations, including larval sleep bout number (Duboué et al. 2011), daily sleep duration (Yoshizawa et al. 2015), adult bout duration and number of bouts per day (Jaggard et al. 2017), and rebound after deprivation (McGaugh et al. 2020). Likewise, 3-week old Tinaja cavefish consume more black worms during a single feeding trial than surface fish, but 3-week old Pachón cavefish consumes less than surface fish (Aspiras et al. 2015). Pachón cavefish also begin fat deposition earlier in development than the other cavefish populations and surface fish (Xiong et al. 2022).

We hypothesize that differences in repeatability among traits are in part caused by differences among cave environments. For example, perpetual darkness is a uniformly similar environmental factor across all caves considered here. Of the traits we examined, the visual sensory system in *A. mexicanus* is likely the most uniformly impacted across independent populations because some light is needed to have vision (Nilsson 2009; Oakley and Speiser 2015). Eye loss appears to exhibit the most uniform genetic basis across independent cave lineages as well. While perpetual darkness disrupts circadian rhythms as well (with cascading impacts on sleep and metabolism (Beale et al. 2013; Moran et al. 2014; Mack et al. 2021; Olsen et al. 2023; Keene et al. 2024)), an organism’s circadian rhythms can be entrained to some extent by other factors such as food intake patterns and interactions with other organisms (Sharma and Chandrashekaran 2005; Abraham et al. 2013; Lewis et al. 2020). Variation in co-existing fauna, including those that enter and leave the caves cyclically like bats (Elliot 2018), among caves likely provide a source of variation for circadian cues. Diets seem to vary among cave populations as well (Espinasa et al. 2017; Wilson et al. 2021; Hernández-Lozano et al. 2025) likely influenced by hydrological variation in caves that impacts water chemistry, flooding frequency, groundwater influx (Elliott 2018; Bauhus et al. 2025; Legendre et al. 2025). The lower genetic repeatability of sleep and metabolic traits may reflect the variation among caves in access to alternative sources of circadian cues and food, allowing more evolutionary paths for these traits.

Despite this phenotypic variation, we observe instances of genetic repeatability across three of the four traits we examined: feeding, sleep, and eye phenotypes. These findings indicate that, even when phenotypes differ in magnitude, direction, or expression timing, evolution may repeatedly target similar chromosomal neighborhoods, suggesting shared genomic constraints or accessible evolutionary routes.

### Repeated use of the same genetic regions for cave trait evolution

The repeated involvement of the same genomic regions across independent cave lineages aligns with theoretical predictions that genetic architectures involving large-effect loci underlying locally favorable traits are easier to maintain when spatially divergent selection occurs in the face of gene flow (Yeaman and Otto 2011b; Yeaman and Whitlock 2011). This could lead to selection repeatedly maintaining particular genetic architectures with larger effects on locally adaptive traits (Battlay et al. 2024) despite there likely being many possible genetic architectures for the traits (Boyle et al. 2017; Liu et al. 2019). Consequently, independent populations could end up using the same large-effect genetic architectures when they are similarly challenged by high migration of non-locally adapted alleles or strong genetic drift (Yeaman et al. 2018; Battlay et al. 2024). The genetic repeatability we observed in our study may in part be the result of similar gene flow histories among independent lineages, as both Lineage 1 and Lineage 2 cave populations have either experienced or continue to experience introgression from surface fish populations (Herman et al. 2018; Moran et al. 2023).

Under local adaptation in the face of gene flow, favored genetic architectures could involve single mutations in genes with large impacts on one or more traits or a locally favorable combination of mutations with a range of effect sizes tightly-linked together in a region of the genome (Yeaman 2013; Yeaman et al. 2016). Some of the most iconic cases of repeated evolution involve mutations in the same gene across lineages, suggesting that repeated adaptation is biased towards certain genes with large mutational targets, standing variation, or pleiotropic roles (e.g. (Conte et al. 2012; Yeaman et al. 2018)). However, we find that genetic variation at higher levels of genome organization may be driving the repeated use of the same genomic regions in cavefish. Only 0.9% (30 of 3,331) of the genes inside the QTL regions we identified have undergone repeated bouts of strong selection across both independent cave lineages. In contrast, 6% (214 of the 3,331) of genes inside QTL regions showed evidence of sweeps in individual cave populations. The paucity of shared signatures of selection in genes could stem in part from unknown differences in demography and selection intensity among cave populations leading to unequal detection power for detecting selection. However, our previous study using a similar selection test found that approximately 30% of selective sweeps detected within one cave population can also be detected in at least one cave population from the other cave lineage (Moran et al. 2023), which far exceeds the percentage of genes we found shared signatures of selection for in our QTL regions.

We hypothesize that the genetic repeatability underlying the repeated loss of sleep and eyes among cave populations may be driven by the clustering of functionally related loci, with a range of individual effect sizes on our focal traits, rather than repeated evolution for mutations in the same set of single genes that selection could act strongly upon individually.

### Genetic co-localization among cave traits is both trait- and population-dependent

While repeated use of the same genomic regions across independent cave populations was a present in the genetic basis of all repeated cave traits we examined, genomic co-localization among the genetic bases of these traits was rare. We detected only two instances of QTL co-localization among traits: a region chromosome 5 linking sleep and eye size and a region on chromosome 25 linking sleep and feeding. Our QTL mapping suggests that the evolution of sleep and feeding amount behaviors in cavefish has been largely decoupled, despite functional interactions between these traits being observed across animals (Stahl et al. 1983; Murphy et al. 2016; Makino et al. 2021; Gallman et al. 2024).

While uncommon among the traits we examined, the genetic co-localization of several QTL may still explain some of the correlated phenotypic evolution we observe within a cave population. For the Surface x Tinaja F2 mapping population, sleep and feeding QTL mapped to the same region of chromosome 25. This region of chromosome 25 houses two promising candidate genes (*acsl1a* and *mtnr1aa*) with known effects on sleep and eating behavior. *Ascl1a* is an ortholog of the mammalian long-chain fatty acyl-CoA synthetase 1 (*ascl1*) gene that plays a key role in fatty acid metabolism. *Ascl1* has been associated with metabolic traits in model organisms like insulin resistance, but also dysregulated circadian rhythms (Osipovich et al. 2023; Peng et al. 2024; Zheng et al. 2025). In *A. mexicanus*, *acsl1a* is under selection in wild populations from Lineage 2 only: Pachón cave, Tinaja cave and Rascón surface population. Interestingly, *acsl1a* is also significantly upregulated in the livers of both Tinaja and Pachón individuals compared to Río Choy surface individuals and associated with a cave-specific open chromatin region of a cis-regulatory element nearby the gene (Krishnan et al. 2022). *Mtnr1aa* is a homolog for the mammalian gene *mntr1* that encodes the MT1 receptor protein for melatonin. Melatonin and its receptors are best known for their involvement in sleep/wake cycles, but they can also influence glucose homeostasis, including insulin resistance (Contreras-Alcantara et al. 2010; Owino et al. 2018; Buonfiglio et al. 2019). In *A. mexicanus*, *mtnr1aa* is not uniquely under selection in cave environments, as selective sweeps are observed in all three cave and both surface wild populations we examined (Supplementary Data S4). However, Tinaja cavefish uniquely appear to have lost endogenous melatonin rhythms altogether; they have low levels of melatonin throughout the entire day even under light:dark conditions (Mack et al. 2021). While Molino and Pachón cavefish do display lower melatonin levels under dark:dark conditions, they still produce surface-like melatonin rhythms with lower levels during the day and higher at night under light:dark cycles (Mack et al. 2021). The Tinaja-unique QTL for sleep and feeding on chromosome 25 suggests that unique mutations or expression patterns in *mtnr1aa* may help explain the nuances observed in circadian rhythm loss among caves.

The sparse examples we found for genetic co-localization among sleep, feeding and eye size in cavefish stand in contrast to a major synthesis of most of the previous QTL studies done predominantly on Surface x Pachón cavefish mapping populations where genetic co-localization is frequent across the genome (Wiese et al. 2025). None of these previous studies included QTL regions associated with sleep or eating amount, but many have included eye size. Eye size in Surface x Pachón cavefish does appear to co-localize with a variety of cave traits from pigmentation to taste bud and tooth count to body condition (Protas et al. 2008; Yoshizawa et al. 2012; Wiese et al. 2025). The extensive variation in both traits and populations where genetic co-localization occurs in this system provides an interesting opportunity to dissect the types of mutations, trait interactions, and selective regimes that make genetic co-localization among distinct traits more or less likely to occur.

### Candidate genes for the repeated genetic co-localization of sleep and eye size to chromosome 5

Regardless of the generality for genetic co-localization among sleep, eye size and eating behavior across the genome, we can leverage the one instance of co-localization that’s repeated among lineages to investigate whether repeated, pleiotropic mutations in single genes drive this signal. The QTL region on chromosome 5 associated with sleep in all three hybrid populations is also associated with eye size in the Surface x Molino F2 hybrid population (Table 1 & 2). While we did not detect an association between Lineage 2 cave ancestry and eye size in this region with our dataset, eye size QTL have been detected in this region in previous Surface x Pachón mapping populations as well (Wiese et al. 2025). This region of chromosome 5 contains several genes with known pleiotropic effects relevant for cave trait evolution: *tgfa*, *oca2*, and *ghrl*. We will discuss each gene and the evidence for their potential pleiotropic impacts on *A. mexicanus* cave phenotypes in turn.

*Tgfa*, which we established a role for in sleep outcomes in this study, plays a role in retinal cell differentiation during eye development (Luetteke et al. 1993; Reneker et al. 1995). To the best of our knowledge, none of the mutations introduced to *tgfa* so far have significantly modified both eye development and sleep simultaneously within one organism. *Tgfa* has been experimentally linked to eye development through null *tgfa* mutant mice with reduced eye size (Luetteke et al. 1993), and sleep through null *tgfa* mutant zebrafish with reduced sleep (Lee et al. 2019). The *tgfa* surface *crispants* in our study also slept significantly less than their wild-type siblings, further support a major role for *tgfa* in modifying sleep patterns. *tgfa* crispants were also included in a recent eye screen for gene candidates involved in cavefish eye loss (Shennard et al. preprint). *Tgfa* crispants exhibited subtly smaller eye and pupil sizes and worse optomotor response compared to wild-type counterparts. However, these eye effects were non-significant after false discovery rate corrections, suggesting the introduced mutations in *tgfa* did not reduce eye size and sleep to equivalent degrees. *tgfa* is downregulated in the degenerating eyes of 54 hpf Pachón cavefish relative to age-matched surface fish (Gore et al. 2018), so it is possible that regulatory changes to *tgfa* during eye development also play a role in eye loss in cavefish. No cave populations have experienced strong selective sweeps on intragenic variation within *tgfa,* but one surface populations has experienced a selective sweep in this gene as well (Supplementary Data S4). Together, this suggests that *tgfa* may potentially contribute to eye and sleep phenotypes observed in cavefish and is worthy of future in-depth studies.

This multi-trait QTL region contains another multi-functioning gene*, ghrl,* which encodes the peptide hormone Ghrelin. On paper, ghrelin is promising candidate for inducing simultaneous changes in sleep, appetite, and eye development in *A. mexicanus* given well-established roles in appetite stimulation, wakefulness and neuroprotection against several eye diseases in mammals (Bodosi et al. 2004; Steiger 2007; Zaniolo et al. 2011; Azevedo-Pinto et al. 2013; Bai et al. 2021). For *A. mexicanus* however, we did not find an association between this QTL region housing ghrelin and eating behavior. This result aligns with a recent finding that ghrelin signaling does not drive appetite in zebrafish in the same way it drives appetite in mammalian model organisms (Opazo et al. 2019; Ahi et al. 2022). *Ghrl* is also not among the genes differentially expressed in the degenerating eye tissue of 54 hpf Pachón larval fish compared to age-matched surface fish (Gore et al. 2018). Cavefish population also did not experience strong selective sweeps on intragenic variation within *ghrl* (Supplementary Data S4). While we do not yet have direct functional evidence for the impact of *ghrl* on cave traits; the indirect lines of evidence available to us do not support a pleiotropic role for ghrelin influencing sleep loss, eye loss, and eating behavior simultaneously.

The other promising gene candidate for inducing simultaneous changes in multiple traits in this region of chromosome 5 is the gene *oca2.* Unlike *tgfa and ghrl,* there is functional evidence that a loss-of-function mutation in *oca2* can cause simultaneous shifts to multiple cave traits. Surface fish homozygous for a null *oca2* mutation sleep less, lack pigment, and have a reduced stress response compared to their wildtype siblings. Null *oca2* fish however, do not have significantly smaller eyes than wildtype counterparts (Klaassen et al. 2018; O’Gorman et al. 2021; Choy et al. 2025a). *oca2* has also not been associated with gross eye size differences in other organisms (Kamaraj and Purohit 2014; Neveu et al. 2022). Therefore, while loss-of-function mutations in *oca2* have likely played a simultaneous role in sleep loss and other cave traits like albinism, they likely do not contribute directly to the reduction of eye size in *A. mexicanus*.

In sum, the current lines of genetic evidence we have in *A. mexicanus* support a role for linkage between multiple variants with effects on individual traits. Pleiotropic mutations in this region may have also contributed to the repeated use of this genomic region, but do not appear to be a strong constraining factor across populations through repeated, strong selection for them.

### The utility of QTL studies in Astyanax for understanding the genetic basis of complex traits

We anticipate that the candidate genes and QTL regions identified in this study will serve as valuable targets for future functional and comparative analyses. Three CRISPR-Cas9 functionally validated genes capable of modifying sleep in *A. mexicanus* lie within the QTL regions we detected, which represent natural genetic variation underlying divergent sleep between cave and surface fish. We have already discussed two of these genes, *tgfa* and *oca2*. The third gene functionally validated for sleep in *A*. *mexicanus* is *rorca*, a circadian clock gene. Surface fish injected with Cas9 and a gRNA targeting *rorca* slept less on average at night compared to their wild type siblings (Mack et al. 2021). The presence of a nighttime sleep QTL in the region of chromosome 16 that houses *rorca* and a previous finding that cavefish populations from both lineages have lost rhythmic expression of *rorca* (Mack et al. 2021) again highlight the discovery of natural genetic variation in *rorca* influencing sleep. Though we only highlighted a few other candidate genes in text (e.g. *ube3a*, *acsl1a*, *mtnr1aa*), many other candidate genes identified and validated in other model organisms fall inside the QTL regions of this study (Supplementary Data S4). The QTL maps presented here (which represent QTLs with the highest marker density and highest sample size in *A. mexicanus* to date), combined with our synthesis of previous QTL studies (Tables S29–S31), expression data, and selective sweep analyses (Tables S21–S25, Supplementary Data S4), provide a valuable framework for both validating whether genes identified in lab-based studies contribute to natural variation in complex traits like sleep and for identifying novel candidates for future functional validation and mechanistic work.

### Conclusion

We discovered extensive genetic repeatability across independent cavefish lineages through our approach of extending QTL analysis to multiple populations. However, the repeated co-evolution of eye loss, sleep loss, and metabolic shifts in appetite and fat deposition does not appear to be driven by wide-spread genetic correlation between these traits. We found little evidence for the repeated use of specific genes driving the repeated evolution of these traits but found evidence for use of the same genomic regions across independent lineages. This study provides a foundation for exploring features of the *A. mexicanus* genome that constrain evolution to repeated outcomes. Additionally, the uniquely distinct divergence in sleep patterns between interbreeding ecotypes of the *A. mexicanus* provided us with an unparalleled opportunity to map the genetic basis of sleep in a natural system for the first time. We found that several previously proposed candidate genes from functional experiments such as *oca2*, *rorca*, and *tgfa* are likely involved in the natural evolution of sleep loss and are genetically correlated to other traits. However, colocalizing genetic architectures among traits are not strongly repeated across replicate cave populations, which suggests that genetic repeatability through genetic correlation among coevolving traits is not a highly predictable outcome in repeated evolution overall.

## Data availability

The reference genomes used in this study can be found through NCBI at the following accession: GCA_023375975.1. The RAD-seq raw sequence data for the mapping populations can be obtained through NCBI BioProject PRJNA1282214 and NCBI SRA accessions for the WGS of the wild parent populations can be found in Supplementary Data S7. The genotype and phenotype data used as input for the QTL analysis can be found in Supplementary Data S1 and S2 files. Scripts associated with the QTL analyses and overlap of QTL regions with other genome features are in the following repository: https://github.com/emiliejrichards/Astyanax-MultiLineage-Sleep-Feeding-QTL/ Scripts associated with quantifiying selective sweeps across the genome are in the following repository: https://github.com/robackem/Cavefish-SV

## Acknowledgments

We would like to thank the Minnesota Supercomputing Institute (University of Minnesota) for access to the high-performance computing systems that enabled this computationally intensive research project. We also thank Maya Enriquez, Danielle Drabeck, Manisha Munasinghe, Sam Yeaman and the Yeaman Lab for providing feedback on early drafts of the manuscripts.

## Study funding

This work was supported by National Institutes of Health grant R01GM127872 awarded to S. McGaugh, N. Rohner, and A. Keene, National Institutes of Health grant R35GM138345 to J. Kowalko, and National Science Foundation grant DEB 2316783/2316784 to S. McGaugh and J. Kowalko. The R01GM127872, DEB-2316783, and Minnesota IRACDA grants supported E. Richards during the preparation of this manuscript.

## Conflict of interests

All authors declare that they have no known competing financial or personal interests that would influence the work shown in this manuscript.

